# Phanerozoic Radiation of Ammonia Oxidizing Bacteria

**DOI:** 10.1101/655399

**Authors:** LM Ward, DT Johnston, PM Shih

## Abstract

The modern nitrogen cycle consists of a web of microbially mediated redox transformations. Among the most crucial reactions in this cycle is the oxidation of ammonia to nitrite, an obligately aerobic process performed by a limited number of lineages of bacteria (AOB) and archaea (AOA). As this process has an absolute requirement for O_2_, the timing of its evolution – especially as it relates to the Great Oxygenation Event ~2.3 billion years ago – remains contested and is pivotal to our understanding of nutrient cycles. To estimate the antiquity of bacterial ammonia oxidation, we performed phylogenetic and molecular clock analyses of AOB. Surprisingly, bacterial ammonia oxidation appears quite young, with crown group clades having originated during Neoproterozoic time (or later) with major radiations occurring during Paleozoic time. These results place the evolution of AOB broadly coincident with the pervasive oxygenation of the deep ocean. The late evolution AOB challenges earlier interpretations of the ancient nitrogen isotope record, predicts a more substantial role for AOA during Precambrian time, and may have implications for understanding of the size and structure of the biogeochemical nitrogen cycle through geologic time.

## Introduction

The biogeochemical nitrogen cycle is second only to carbon in size and, arguably, importance for the biosphere (e.g. Tyrrell 1999). The nitrogen cycle supplies fixed nitrogen for biomass while also fueling diverse microbial metabolisms, with fluxes hundreds of teramoles of nitrogen per year (e.g. Canfield et al. 2010). Nitrogen primarily enters this cycle by way of reduced forms (i.e. ammonia fixed from N_2_ by the enzyme nitrogenase), and so biological nitrification (i.e. the oxidation of ammonia to nitrite and nitrate) is an essential step for enabling downstream processes such as anammox and denitrification (Zerkle and Mikhail 2017). No metabolism has yet been discovered that is capable of oxidizing ammonia in the absence of O_2_ or O_2_-derived compounds like nitrite or NO (Hu et al. 2019)—therefore the modern nitrogen cycle where oxidized forms are regenerated and recycled are necessarily tied to O_2_.

Aerobic oxidation is found in a limited, polyphyletic set of Bacteria (AOB) and Archaea (AOA). In both AOB and AOA, the first step in ammonia oxidation is performed via ammonia monooxygenase (AMO), a member of the copper membrane monooxygenase (CuMMO) family. The CuMMO family includes the related particulate methane monooxygenases (pMMO) and enzymes that oxidize other small hydrocarbons (Khadka et al. 2018). CuMMO enzymes have an absolute requirement for O_2_, leading to the hypothesis that metabolic pathways utilizing these enzymes—including ammonia oxidation—evolved after the evolution of oxygenic photosynthesis provided significant O_2_ to the environment. While alternative, O_2_-independent ammonia oxidation processes such as the coupling of ammonia oxidation to phototrophy or metal reduction have been hypothesized, no organism has ever been characterized that can perform these reactions (Ward et al. 2019b, in ‘t Zandt et al. 2018). The evolution of oxygenic photosynthesis in Cyanobacteria led to the accumulation of atmospheric O_2_ to biologically meaningful concentrations ~2.3 billion years ago (Ga) during the Great Oxygenation Event (GOE), and it has been suggested that the onset of the aerobic nitrogen cycle occurred shortly thereafter (Zerkle et al. 2017). However, others have argued from isotopic evidence that an aerobic nitrogen cycle was in place much deeper in Earth history (e.g. Garvin et al. 2009). Distinguishing between these possibilities from the rock record alone is difficult due to the poor preservation of Archean strata and the lack of a robust framework for interpreting the ancient nitrogen isotope record. Moreover, signatures in the nitrogen isotope record may reflect only the expansion to geochemical prominence or first preservation of signatures of nitrogen metabolisms, and not necessarily their initial evolutionary origin. Instead, the biological record can provide opportunities for querying the antiquity of organisms and metabolisms responsible for driving the nitrogen cycle.

Here, we estimate the antiquity of AOB via phylogenetic and molecular clock analyses. We show that ammonia oxidation in bacteria has evolved convergently at least twice and that crown group AOB clades originated <1 Ga, with major radiations occurring within the last ~500 million years (Ma). This suggests that bacteria did not continue to ammonia oxidation until late in Earth history—more than 1.5 Ga after O_2_ first accumulated in the atmosphere. The predicted appearance of AOB at a time when Earth surface environments underwent oxygenation to modern-like levels points to the potential role for niche expansion in fostering evolution and boosting turnover of the marine fixed nitrogen inventory. This suggests a substantial difference in scale or structure of the biogeochemical nitrogen cycle during Precambrian time, likely with a more dominant role for AOA.

### Phylogenetic distribution of proteins involved in ammonia oxidation

Phylogenetic analysis of the distribution of genes associated with ammonia oxidation shows that this metabolism is restricted to four clades of characterized ammonia oxidizers (Figure 1). These include members of the Nitrosphaeria class of Crenarchaeota/Thaumarchaeota, the Nitrococcaceae family within the Gammaproteobacteria, the Nitrosomonadaceae family within the Betaproteobacteria, and some members of *Nitrospira* (the “comammox” bacteria, the only known organisms capable of oxidizing ammonia to nitrite and subsequently to nitrate, van Kessel et al., 2015, Daims et al., 2015)(Figure 2). These results are based on ammonia monooxygenase and homologous proteins from the copper membrane monooxygenase family (Figure 2, Supplemental Figures 1 and 2). While CuMMO sequences were recovered from diverse lineages including some that have not previously been characterized to possess the capacity for methanotrophy (e.g. members of the UBP10 and Myxococcota phyla), these proteins are most closely related to enzymes that are characterized as performing carbon oxidation (e.g. pMMO, butane monooxygenase), and no organisms outside of characterized clades of ammonia oxidizers were found to encode AMO. In all cases, ammonia oxidation appears to be a derived trait, with basal members of the clades and closely related outgroups lacking the capacity for ammonia oxidation. Importantly, these clades of ammonia oxidizing microorganisms are not closely related and are phylogenetically separated by many lineages incapable of ammonia oxidation (Figure 1).

**Figure 1:**
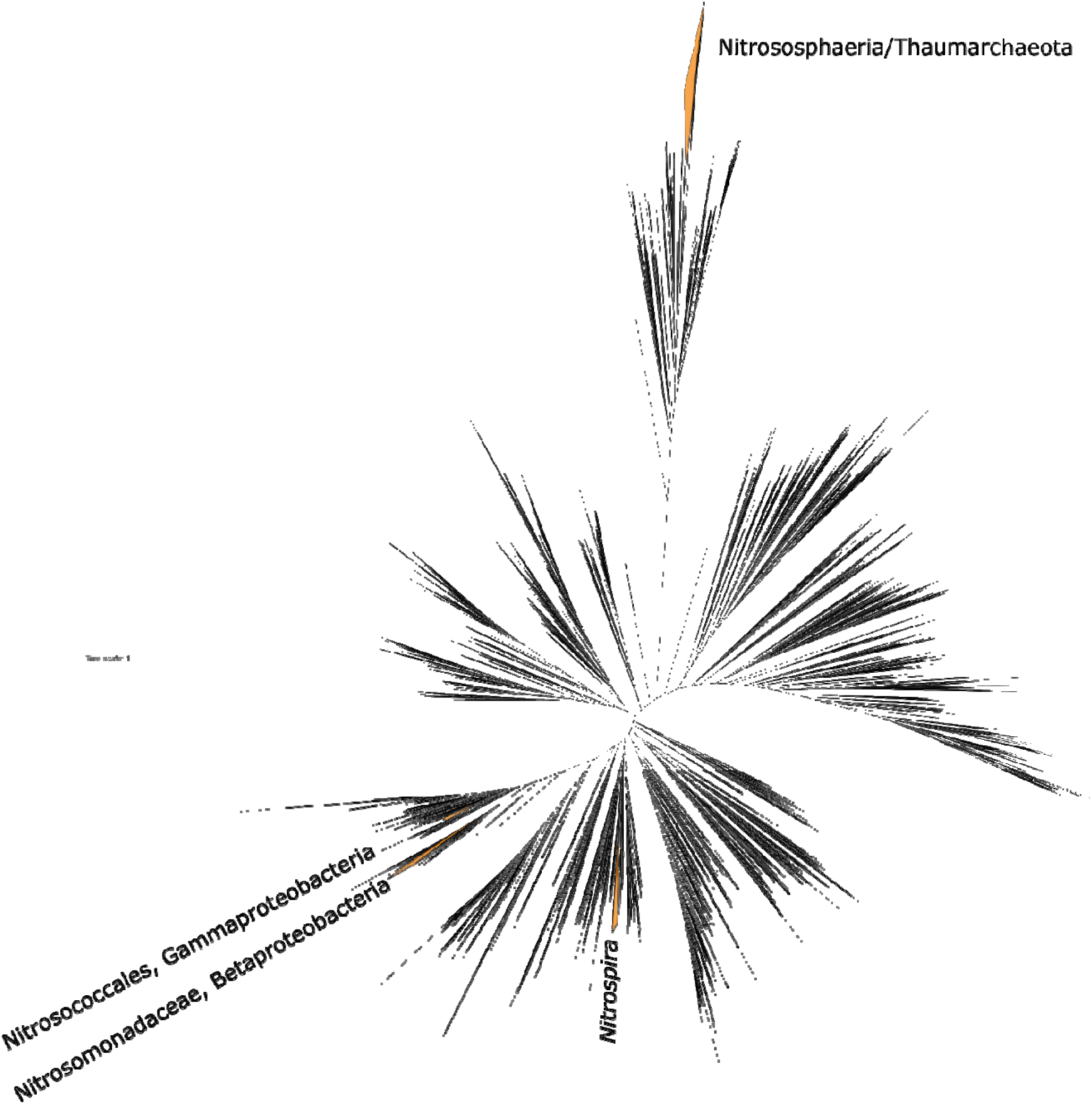
Tree of life built with concatenated ribosomal proteins following methods from Hug et al. 2016. Clades of ammonia oxidizing organisms highlighted in orange and labeled. The distribution of ammonia oxidation is polyphyletic, spread across one lineage within the Archaea (Nitrososphaeria) and three within the Bacteria (Nitrosococcales and Nitrosomonadaceae in the Proteobacteria phylum, and some members of the genus *Nitrospira* within the Nitrospirota phylum).

**Figure 2:**
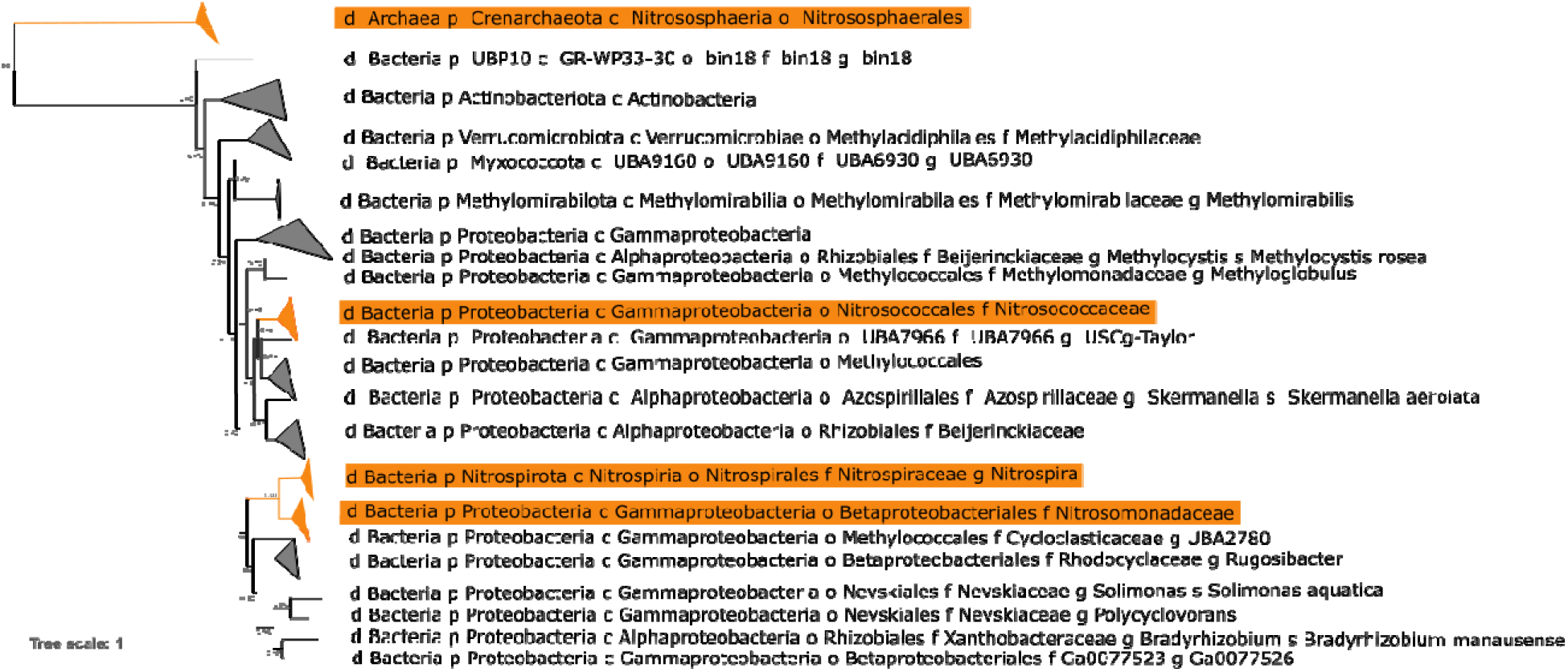
Phylogeny of concatenated protein sequences of CuMMO A and B subunits (i.e. AmoA and AmoB, PmoA and PmoB), with ammonia monoxygenases highlighted in orange. Major clades are collapsed and labeled by the taxonomy of the organisms in which they are found as determined with GTDB-Tk (Parks et al. 2018). The highly divergent archaeal ammonia monooxygenase is placed as an outgroup, though the placement of the root is indeterminate. Transfer Bootstrap Expectation (TBE) support values shown as calculated by BOOSTER (Lemoine et al. 2018).

Phylogenetic analysis of proteins involved in ammonia oxidation, compared to organismal relationships among AOB, provide evidence for convergent evolution and horizontal gene transfer as major drivers for the extant diversity of organisms with the genetic capacity for ammonia oxidation. These relationships are consistent with major clades of AOB acquiring the capacity for ammonia oxidation through separate evolutionary events, followed largely by vertical inheritance within each AOB clade. These data are not consistent with a much more ancient acquisition of ammonia oxidation (e.g. in the last common ancestor of Nitrosococcales and Nitrosomonadaceae) followed by extensive loss. As a result, the age of total group Nitrosococcales, Nitrosomonadaceae, and the extant diversity of comammox *Nitrospira* can confidently be used to constrain the timing of acquisition of the capacity for ammonia oxidation in each lineage. Additionally, our data are consistent with hypotheses for ammonia oxidation evolving from earlier aerobic methane oxidation pathways (Supplemental Information).

### Molecular clock evidence for the late evolution of ammonia oxidizing bacteria

To connect the evolution history of AOB described above to events in Earth history, we performed molecular clock analyses to determine when AOB clades diverged from non-ammonia oxidizing relatives (i.e. age of total groups) and when AOB clades subsequently radiated (i.e. age of crown groups). Molecular clocks estimate the origin of each AOB clade to Neoproterozoic or Phanerozoic time, with each stem group AOB lineage emerging between 238 (comammox *Nitrospira*) and 894 Ma (Nitrosococcaceae) and radiation of crown groups occurring after 538 Ma (Figure 3, Table 1). The 95% confidence intervals of divergence times introduce uncertainty of +/- 150 Ma to these estimates (Table 1, Supplemental Figure 4), but in all cases firmly place the major radiation of extant AOB to Phanerozoic time even if the origin of crown groups was in late Neoproterozoic time. Between the uncertainty in the ages of total group AOB clades, and the fact that the acquisition of ammonia oxidation could in theory occur at any point along stem lineages prior to the divergence of crown groups, the range of 95% confidence intervals of ages of origin of the first AOB consistent with our data is between 1169 and 414 Ma. This age range includes scenarios involving the first evolution of ammonia oxidation in the earliest stem group Nitrosococcaceae (1169 Ma) or at the base of crown group Nitrosomonadaceae (414 Ma). This also accommodates the possibility that these groups acquired ammonia oxidation roughly simultaneously between 414 and 490 Ma. The analysis therefore does not allow for a unique determination of which proteobacterial lineage first acquired the capacity for ammonia oxidation. However, all scenarios consistent with our data involve a later acquisition of ammonia oxidation within the *Nitrospira*, after the radiation of ammonia oxidizing Nitrosomonadaceae and nitrite oxidizing *Nitrospira*.

**Figure 3:**
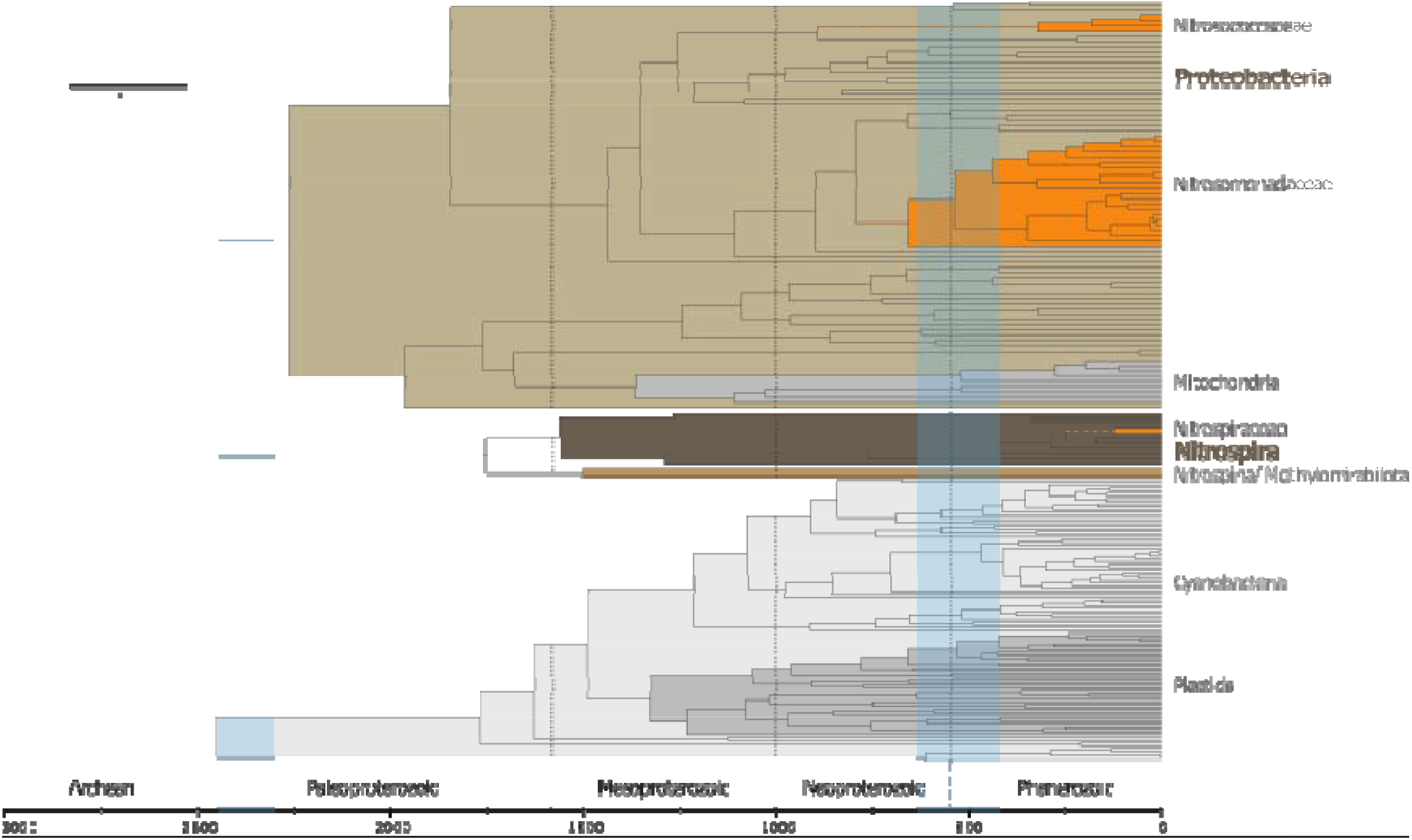
Molecular clock showing estimated age of clades of ammonia oxidizing bacteria. Phylum-level clades highlighted in gray and brown, with ammonia oxidizing clades highlighted in orange. Approximate timing of Great Oxygenation Event (~2.45-2.3 Ga) and Neoproterozoic/Paleozoic Oxygenation Event (~635-420 Ma) shown with light blue bars. While stem lineages of AOB clades may predate the NOE, the radiation of crown groups all occur broadly coincident or subsequent to the NOE, suggesting that evolutionary radiations of nitrogen-cycling organisms may have been causally linked with expansions in ocean oxygenation and/or productivity during this time period.

**Table 1:** Age ranges for key divergences discussed in the text in millions of years (Ma).

| Clade | Estimated age of crown group | 95% Confidence Interval | Estimated age of total group | 95% confidence interval |
| --- | --- | --- | --- | --- |
| **Nitrosococcaceae** | 325 | 194-490 | 894 | 596-1169 |
| **Nitrosomonadaceae** | 538 | 414-662 | 657 | 510-821 |
| **Nitrospira** | 124 | 59-197 | 238 | 149-329 |

The Neoproterozoic to early Phanerozoic origin of crown group AOB suggested by our data is ~1.5 Ga later than previous suggestions that placed ammonium oxidation at or before the GOE ~2.3 Ga (e.g. Garvin et al. 2009, Zerkle et al. 2017), suggesting that the first rock record evidence for ammonia oxidation records the activity of ammonia oxidizing archaea or other biological or abiotic processes. Our estimate for the origin of bacterial ammonia oxidation is during the time in Earth history that saw the biosphere transition from a low-productivity, exclusively microbial state characteristic of Proterozoic time (Dick et al. 2018, Ward and Shih 2019) to a more modern system fueled by eukaryotic algae and supporting complex multicellular organisms including animals (Erwin et al. 2011, Brocks et al. 2017) possibly triggered by increased phosphate availability (Laakso et al. 2020). Net primary productivity of the biosphere is thought to have increased significantly at this time (Ward et al. 2019a, Ward and Shih 2019), along with a rise in atmospheric oxygen concentrations to near-modern levels and the more permanent oxygenation of the deep ocean (Sperling et al. 2015). The timing of this final rise in atmospheric and marine O_2_ is not well-constrained and may have occurred as early as Ediacaran time (Lyons et al 2014) or as late as ~420 Ma (e.g. Stolper and Keller 2018). The divergence of stem group proteobacterial AOB during Neoproterozoic time and the radiation of crown group proteobacterial AOB clades during Paleozoic time suggests that these evolutionary innovations may be causally linked. Increased oxygenation of the oceans would have provided additional O_2_ for ammonia oxidation, while higher NPP necessitates higher fluxes of fixed nitrogen through the biosphere leading to higher rates of N_2_ fixation to reduced forms (e.g. Ward et al. 2019a). As a result, the necessary substrates for ammonia oxidation would have been more abundant after this later rise of O_2_ than earlier in Proterozoic time, potentially opening additional niche space that enabled the radiation of AOB. This hypothesis is further supported by the tendency of AOB to be adapted to higher ammonia (e.g. Martens-Habbena et al. 2009, Schleper 2010, Kits et al., 2017, Hink et al. 2018) and possibly oxygen (e.g. Ke et al. 2015, Qin et al. 2017) concentrations than AOA.

The delayed evolution and radiation of AOB may also be a consequence of limited copper availability in Proterozoic oceans. Aerobic ammonia oxidation has a relatively high requirement for copper for enzyme cofactors (e.g. Amin et al. 2013), and copper availability in Proterozoic oceans may have been limited due to the insolubility of copper sulfide minerals in periodically euxinic oceans (e.g. Saito et al. 2003) and/or lower continental weathering of copper and subsequent runoff into the oceans (Hao et al. 2017). The expansion of some metabolic pathways may therefore have been impeded by the availability of trace metals necessary as enzyme cofactors (Saito et al. 2003). AOA have a comparable copper requirement for electron transport and nitrogen metabolic proteins as AOB (e.g. Walker et al., 2010), and so extreme copper limitation during Proterozoic time would be expected to impede AOA as well as AOB. Copper limitation may therefore have limited the expansion and potential productivity of the oxidative nitrogen cycle for much of Proterozoic time.

### Implications for the Proterozoic nitrogen cycle

The late evolution of crown group AOB clades suggests that bacteria were not playing a dominant role in driving the oxidative arm of the nitrogen cycle during Proterozoic time, though it is always possible that there is a deeper history of bacterial ammonia oxidation by other lineages that remain undiscovered or that are now extinct. The nitrogen isotope record is consistent with active nitrification and denitrification through most of Proterozoic time (e.g. Stüeken et al., 2016, Koehler et al., 2017, Kipp et al., 2018), but does not provide direct evidence for the taxonomic affinity of organisms driving these processes. While the lack of an archaeal fossil record for calibrating molecular clocks makes estimating the antiquity of AOA challenging, recent work (Ren et al., 2019) has suggested that ammonia oxidizing Thaumarchaeota originated ~2.3 Ga, in time to drive the aerobic nitrogen cycle shortly after the GOE ~2.3 Ga (e.g. Zerkle et al. 2017). Consistent with this hypothesis is the lower oxygen requirements of AOA compared to proteobacterial ammonia oxidizers, leading to the continued dominance of AOA in modern oxygen minimum zones (e.g. Bristow et al. 2016). This is also in keeping with AOA having evolved at a time with one to several orders of magnitude lower O_2_ than was present during the origin of AOB (e.g. Lyons et al. 2014). The relative contribution of AOA and AOB to ammonia oxidation fluxes through time has not previously been constrained as the nitrogen isotope signatures of these groups overlap (Santoro and Casciotti 2011) and the relative abundance and activity of AOA and AOB in modern environments is only determined roughly on a local scale via sequencing-based approaches that are not applicable to deep time (e.g. Bristow et al. 2016). However, our results in combination with those of Ren et al., suggest that the AOA were responsible for driving all biological ammonia oxidation for most of Proterozoic time until the origin of the first AOB <1 Ga.

It is important to note that AOA and AOB utilize different biochemical pathways downstream of AMO, so their relative contribution to ammonia oxidation through time has significant implications for modeling of the productivity and atmospheric impact of the ancient biosphere. For example, extant AOA fix carbon using a uniquely energy-efficient O_2_-tolerant carbon fixation pathway (a variant of the hydroxypropionate/hydroxybutyrate pathway), while AOB typically utilize the Calvin Cycle (or, in comammox *Nitrospira*, the rTCA cycle, Lücker et al. 2010) (Könneke et al. 2014, Ward and Shih 2019). This allows AOA to fix 1.3g of dry cell mass for every mole of ammonia oxidized, in contrast to only 0.8g/mol in ammonia oxidizing Proteobacteria (Könneke et al. 2014). Further, nitrification currently accounts for ~75% of non-photosynthetic carbon fixation in aquatic environments (Raven 2009), so a nearly twofold difference in efficiency of carbon fixation in AOA versus AOB may lead to significant differences in predictions of net primary productivity of the biosphere through time. This is particularly important in the Proterozoic when photosynthetic carbon fixation rates are thought to have been much lower than today (e.g. Crockford et al. 2018, Ward and Shih 2019, Hodgskiss et al., 2019). Furthermore, the typical release of N_2_O by AOA is significantly lower than from AOB (Hink et al. 2018), particularly under low-oxygen conditions (Stieglmeier et al. 2014). As a result, an increased contribution of AOA to nitrification during Proterozoic time would likely be associated with a lower biogenic N_2_O flux, potentially at levels sufficiently low to prevent N_2_O from accumulating as an important greenhouse gas in the Proterozoic atmosphere as previously proposed (e.g. Buick 2007).

Our results provide necessary constraints for establishing a timeline for the evolution of the biological nitrogen cycle. Before the origin of the first ammonia oxidizers, the nitrogen cycle would have consisted primarily of a vector toward reduced forms, with perhaps some oxidized nitrogen produced abiotically via processes like lightning (e.g. Navarro-Gonzalez et al., 1998, Wong et al. 2017). This reduced biogeochemical nitrogen cycle is thought to have persisted through Archean time (e.g. Ward et al. 2019a, Yang et al. 2019) and may have continued into Proterozoic time until the evolution of the first ammonia oxidizing Archaea. Due to the stability of fixed nitrogen in the oceans as ammonia at this time, the nitrogen demands of phototrophic primary productivity would have been readily met (Ward et al., 2019a, Yang et al., 2019). Following the evolution of the AOA, the Proterozoic biosphere may have still been nitrogen limited, as the conversion of ammonia to nitrite/nitrate in oxygenated surface oceans would likely be followed by substantial loss of fixed nitrogen via denitrification and anammox in anoxic bottom waters (e.g. Fennel et al. 2005). This extensive nitrogen loss would have maintained low concentrations of fixed nitrogen in the oceans, consistent with the relatively high substrate affinity of AOA (e.g. Martens-Habbena et al., 2009) and low overall GPP predictions for that time (Crockford et al., 2018, Ward et al., 2019a, Hodgskiss et al., 2019).

The Earth experienced several evolutionary and environmental revolutions during Neoproterozoic and Paleozoic time including the rise of atmospheric oxygen to near-modern levels (e.g. Lyons et al., 2014, Stolper and Keller 2018), persistent oxygenation of the deep oceans (e.g. Stolper and Bucholz 2019), the rise of eukaryotic algae and animals (e.g. Brocks et al., 2017), and finally the evolution of plants and colonization of terrestrial environments (e.g. Ibarra et al., 2019). These events had a number of effects on weathering and geochemical cycles and may have triggered evolutionary innovations in the nitrogen cycle. For instance, higher primary productivity (e.g. Ward and Shih 2019) would have increased fluxes of nitrogen through the biosphere while increased oxygenation would have increased the stability of nitrate in the oceans and subsequently allowed the accumulation of a large marine fixed nitrogen pool for the first time since the GOE (e.g. Stüeken et al., 2016). These changes may have provided opportunities for the convergent evolution of multiple lineages of ammonia oxidizing bacteria, particularly Nitrosomonadaceae and Nitrosococcaceae.

Finally, it appears that comammox *Nitrospira* evolved last of all known lineages of ammonia oxidizers. Comammox *Nitrospira* appear to be derived from a larger and more ancient clade of nitrite oxidizing Nitrospirota via HGT of ammonia oxidation genes. Molecular clocks suggest that this transition occurred during Mesozoic time (Figure 3). Comammox *Nitrospira* and their nitrite oxidizing relatives are adapted to low O_2_ concentrations (e.g. Palomo et al., 2018); the apparent coincidence of the evolution of comammox *Nitrospira* with Mesozoic Oceanic Anoxic Events (e.g. Robinson et al., 2016) may reflect the expansion of niches for ammonia oxidizers with low oxygen demands at this time.

### Conclusions

The molecular clock evidence for broadly coincident radiations of multiple convergently evolved crown group AOB clades (Nitrosomonadaceae, Nitrosococcaceae, and comammox *Nitrospira*) shown here is largely unprecedented in molecular clock studies, which typically address the age of a single clade (e.g. acquisition of phototrophy within a bacterial phylum, Shih et al. 2017b) or show multiple evolutionary events scattered through time (e.g. evolution of C_30_ sterols in sponges and algae, Gold et al. 2016). This adds strength to interpretations that the convergent evolutionary transitions to ammonia oxidation in these groups and/or their subsequent radiation may be linked to increases in ocean oxygenation at this time and highlights the interconnectedness between evolution of biogeochemically relevant microorganisms and major environmental perturbations. These interpretations are made only stronger in combination with other work indicating evolutionary and ecosystem expansion of AOA around this time (Ren et al., 2019).

The necessity of reevaluating the structure and size of the Proterozoic nitrogen cycle in light of evidence for late-evolving AOB highlights a recurring problem in assessing the antiquity of microbial lineages—the rock record typically records the indirect effects of a (bio)geochemical process or the metabolism driving it, not directly the organisms that perform it. As a result, care must be taken in applying a strictly uniformitarian interpretation of the biological drivers of geochemical processes in deep time, as this overlooks evolutionary processes such as convergent evolution or horizontal gene transfer of metabolic pathways that lead to incongruent histories and potentially different combinations of traits in ancient drivers of biogeochemical cycles from the organisms responsible for these processes today.

## Methods

Phylogenetic methods followed those described previously (Ward and Shih 2020) and summarized here. Genomes were downloaded from the NCBI Genbank and WGS databases. Completeness and contamination of metagenome-assembled genomes (MAGs) was estimated based on presence and copy number of conserved single-copy proteins by CheckM (Parks et al. 2015). Protein sequences used in analyses (see below) were identified locally with the *tblastn* function of BLAST+(Camacho et al. 2009), aligned with MUSCLE (Edgar 2004), and manually curated in Jalview (Waterhouse 2009). Positive BLAST hits were considered to be full length (e.g. >90% the shortest reference sequence from an isolate genome) with *e*-values greater than 1e^-20^. Presence of metabolic pathways of interest in incomplete MAGs was predicted with MetaPOAP (Ward et al. 2018b) to check for False Positives (contamination) or False Negatives (genes present in source genome but not recovered in metagenome-assembled genomes). Phylogenetic trees were calculated using RAxML (Stamatakis 2014) on the Cipres science gateway (Miller et al. 2010). Transfer bootstrap support values were calculated by BOOSTER (Lemoine et al. 2018), and trees were visualized with the Interactive Tree of Life viewer (Letunic and Bork 2016). Taxonomic assignment was confirmed with GTDB-Tk (Parks et al. 2018). Histories of vertical versus horizontal inheritance of metabolic genes was inferred by comparison of organismal and metabolic protein phylogenies (Doolittle 1986, Ward et al. 2018a).

A concatenated protein alignment was generated by extracting protein sequences for marker genes from genomes of interest via the *tblastn* function of BLAST+ (Camacho et al. 2009), aligning protein sequences with MUSCLE (Edgar 2004), and then concatenating aligned sequences. Concatenated alignments were curated with Gblocks (Castresana 2000) and manually in Jalview (Waterhouse 2009). Taxa included in this alignment consist of all available AOB genomes on the NCBI GenBank and WGS databases as well as sister groups and outgroups spanning the full diversity of the Proteobacteria and Nitrospirota as well as closely related phyla (e.g. Methylomirabilota and Nitrospinota) as assessed by GTDB (Parks et al. 2018) and concatenated ribosomal protein phylogenies of the tree of life (Figure 1, Hug et al. 2016), as well as Cyanobacteria, Plastids, and Mitochondria. Phylogenetic markers were chosen as conserved proteins across bacteria, plastids, and mitochondria, as previously reported (Shih et al 2017a, Shih et al. 2017b), and consisted of AtpA, AtpB, EfTu, AtpE, AtpF, AtpH, AtpI, Rpl2, Rpl16, Rps3, and Rps12 protein sequences. Bayesian molecular clock analyses were carried out using BEAST v2.4.5 (Bouckaert et al. 2019) using the Cyberinfrastructure for Phylogenetic Research (CIPRES) Science Gateway v 3.3 server (Miller et al. 2010). As previously reported, the CpREV model was chosen as the best-fitting amino acid substitution model for the concatenated protein dataset based on ProtTest analysis (Shih et al 2017b) Cross-calibration techniques utilizing plastid and mitochondrial endosymbiosis events were used as priors, utilizing time constraints for the most recent common ancestor of Angiosperms (normal distribution with a mean of 217 Ma and sigma of 40 Ma) and of land plants (normal distribution with a mean of 477 Ma and sigma of 70 Ma) as has been previously described by Smith et al (Smith et al. 2010,Shih and Matzke 2013). We also constrained the most recent common ancestor of Rhodophytes with as a more recent study and precise estimate of the fossil constraint Bangiomorpha pubescens utilizing Re-Os isotopic measurements of the Bylot Supergroup of Baffin Island where the fossil was first described (Gibson et al. 2017). Taking into account previously reported ages of Bangiomorpha, we set this constraint as a uniform prior from 1030-1200 Ma, in order to account both Re-Os and Pb-Pb isotopic measurements estimating the age of Bangiomorpha (Butterfield 2000). A conservative uniform prior between 2300-3800 Ma was set on the divergence between Cyanobacteria and Melainabacteria, as oxygenic photosynthesis evolved prior to the Great Oxygenation Event and most likely evolved sometime after the Late Heavy Bombardment. Finally, a uniform prior for all taxa was again set conservatively between 2400-3800 Ma, assuming that the Last Bacterial Common Ancestor most likely evolved after the Late Heavy Bombardment. Wide uniform priors were used as a means to provide very conservative upper and lower limits. Three Markov chain Monte Carlo chains were run for 100 million generations sampling every 10,000^th^ generation, and the first 50% of generations were discarded as burn-in. TreeAnnotator v1.7.5 (Bouckaert et al. 2019) was used to generate maximum clade credibility trees.

As there are no known fossils of archaea to be used as molecular clock calibrations (Ward and Shih 2019), calibrating archaeal molecular clocks with plant and algal fossils requires extrapolating evolutionary rates across the entire Tree of Life. Rates of molecular evolution can vary substantially across deeply diverging lineages (e.g. Kuo and Ochman 2009) and so application of molecular clocks to inter-domain datasets in the absence of robust calibrations can introduce untenable artifacts and uncertainty (e.g. Roger and Hug 2006). Recent molecular clocks spanning the full Tree of Life built with calibrations from only a single domain, for example, can produce dates for the divergence of bacteria and archaea spanning > 4 Ga between different marker sets (Zhu et al., 2019) and with credible intervals spanning >1 Ga for nodes in unconstrained domains (Betts et al., 2018). The methods we utilize here were therefore determined to not be viable for performing molecular clock analyses on ammonia oxidizing archaea and so these organisms were not included in our molecular clock analyses.

## Acknowledgements

LMW acknowledges support from an Agouron Institute Postdoctoral Fellowship, a Simons Foundation Postdoctoral Fellowship in Marine Microbial Ecology, and an NSF XSEDE Startup Award that provided computational resources via the CIPRES Science Gateway.

## Author contributions

All authors conceived the study. LMW collected data and performed phylogenetic analyses. PMS performed molecular clock analyses. LMW and PMS wrote the manuscript with contributions from DTJ.

