## Supplemental Information for "Phanerozoic Radiation of Ammonia Oxidizing Bacteria"

**Supplemental Discussion:**

**Evolutionary relationships of aerobic methanotrophy and ammonia oxidation**

AMO clades show ~59-90% protein sequence similarity across concatenated AmoAB alignments. Curiously, archaeal AMO proteins showed higher sequence-level similarity than bacterial AMO clades (~90% versus ~80% within betaproteobacterial and *Nitrospira* AMO clades, 68% for gammaproteobacterial AMO, and 59% between Betaproteobacteria and *Nitrospira*). In contrast, pMMO clades show somewhat higher amounts of protein sequence-level divergence, with only 52% similarity within the Verrucomicrobiota, 55% similarity among the Methylococcales and UBA7966 Gammaproteobacteria, and 62% similarity within the Beijerinckiaceae.

Ammonia monooxygenase (AMO) is homologous to particulate methane monooxygenase (pMMO) and other proteins in the broader copper monooxygenase (CuMMO) family (Khadka et al. 2018). Phylogenies of the CuMMO family show that AMO sequences are not monophyletic, and instead comprise three separate clades (one comprised of the distantly related archaeal AMO, one comprised of AMO from Gammaproteobacteria and Nitrospirota, and one comprised of AMO from Betaproteobacteria) separated by many lineages of pMMO and other carbon-oxidizing enzymes. These relationships are, broadly, recapitulated in protein phylogenies of hydroxylamine oxidoreductase, responsible for the second step in bacterial ammonia oxidation to nitrite (Supplemental Figure 3). The polyphyly of AMO suggests that proteins in this family have evolved new functions independently multiple times, transitioning between methane- and ammonia-oxidation as previously observed (Khadka et al. 2018). While the phylogeny of AMO and pMMO sequences is unrooted, making confident determination of the directionality of evolution impossible, it seems likely that the ancestral function of this protein family in bacteria was for the oxidation of methane or other hydrocarbons. Clades of AMO sequences have higher sequence identity (are less diverse) than pMMO clades, suggesting that AMO clades are younger, and that the ancestral substrate for this enzyme family was methane. Consistent with this hypothesis is the much broader taxonomic distribution of aerobic methanotrophy relative to ammonia oxidation (e.g. methanotrophy in Verrucomicrobiota, Methylomirabilota, diverse Proteobacteria, and potentially some members of Myxococcota and UBP10, versus ammonia oxidation in only two lineages of Proteobacteria and a small subset of *Nitrospira*)*.* However, if the root is placed on the long branch between bacterial CuMMO sequences and archaeal AMO, it is possible that bacterial pMMO is derived from an ancestral protein that may not have discriminated strongly between methane and ammonia, which is still the case for some members of this enzyme family (e.g. Lontoh et al. 2000).

Evolutionary relationships of CuMMO proteins as well as AOB and aerobic methanotrophs suggest that aerobic methanotrophy may have evolved much earlier than aerobic ammonia oxidation, and that the capacity for ammonia oxidation may have evolved by way of methane oxidation. The sequence-level diversity and taxonomic distribution of pMMO, together with the limited diversity and phylogenetically derived position of AMO, suggests that aerobic methanotrophy had already diversified well before the independent evolutions of bacterial ammonia oxidation, suggesting that bacterial methane oxidation is much more ancient than bacterial ammonia oxidation. This is consistent with interpretations of pMMO arising prior to the divergence between Alphaproteobacteria and Gammaproteobacteria (e.g. Battistuzi et al. 2004), though the distribution of methane monooxygenases is likely a result of ancient horizontal gene transfer events (e.g. Osborne and Haritos 2018). Our data indicates that the origin of aerobic methanotrophy predates that of aerobic ammonia oxidation in bacteria, but constraining the absolute age of this metabolism will require additional investigation. Preliminarily, our data does suggest a post-GOE origin for most extant lineages of aerobic methanotrophic bacteria. Most aerobic methanotrophs are found in the phyla Proteobacteria, Methylomirabilota, and Verrucomicrobia, with some evidence for recent horizontal gene transfer of methanotrophy into other lineages including the Chloroflexi (Ward et al. 2019b). While we do not have an estimate for the age of methanotrophic Verrucomicrobia, the Chloroflexi have previously been shown to have radiated post-GOE (Shih et al. 2017b) and our data here suggest that the radiation of crown group Proteobacteria occurred ~2263 Ma and the divergence of Methylomirabilota from Nitrospinota ~1498 Ma, indicating that methanotrophs in these phyla postdate the GOE.

The shared biochemistry and evolutionary relationships of aerobic ammonia oxidation and methanotrophy, coupled with our estimates for their relative timing of evolution, are consistent with previous hypotheses for the evolution of proteobacterial ammonia oxidation from a methane oxidizing ancestral state (e.g. Khadka et al. 2018). pMMO and AMO show a high degree of crossreactivity between methane and ammonia (Hanson and Hanson 1996), leading to aerobic methanotrophs that incidentally produce a small amount of hydroxylamine from ammonia oxidation (e.g. Bédard and Knowles 1989, Campbell et al. 2011). In order to avoid hydroxylamine toxicity, it is therefore advantageous for aerobic methanotrophs in ammonia-rich environments to encode hydroxylamine oxidoreductase, as is observed in some extant organisms (e.g. Nyerges and Stein 2009, Skennerton et al. 2015). While hydroxylamine oxidation would likely initially have been for the sake of detoxification, electrons from hydroxylamine could be fed into the electron transport chain, making this process metabolically favorable. As a result, incidentally ammonia-oxidizing methanotrophs may have gradually transitioned into incidentally methanotrophic ammonia oxidizers, and eventually evolved into obligate ammonia oxidizing bacteria. This hypothesis is further supported by extant transitional forms, such as ammonia oxidizers that encode methanotrophy genes (Wang et al. 2018) and methanotrophs that encode hydroxylamine detoxification genes (Nyerges and Stein 2009, Skennerton et al. 2015, Ward et al. 2019b). In contrast, it is apparent that ammonia oxidizing *Nitrospira* were ancestrally nitrite oxidiziers that acquired ammonia oxidation in order to drive comammox (Palomo et al. 2018). While previous work has suggested that *Nitrospira* acquired the capacity for ammonia oxidation via horizontal gene transfer from Betaproteobacteria (Palomo et al. 2018), our analyses show *Nitrospira* AMO and HAO genes diverging prior to the radiation of crown group ammonia oxidizing Betaproteobacteria (Figure 2, Supplemental Figures 1-3), suggesting that *Nitrospira* received ammonia oxidation genes from stem group Nitrosomonadaceae or that both groups acquired ammonia oxidation from a third, unknown donor.

**Supplemental figures:**

1. Phylogeny of AmoA/PmoA protein sequences, with TBE support values.
2. Phylogeny of AmoB/PmoB protein sequences, with TBE support values.
3. Phylogeny of Hydroxylamine Oxidoreductase protein sequences, with TBE support values.
4. Full molecular clock, including complete taxon labels and 95% confidence intervals for all nodes.

**Supplemental table:**

1. List of genomes included in molecular clock analysis
