## Supplementary figures and images for "Phanerozoic Radiation of Ammonia Oxidizing Bacteria"

### Supplemental Figure 1

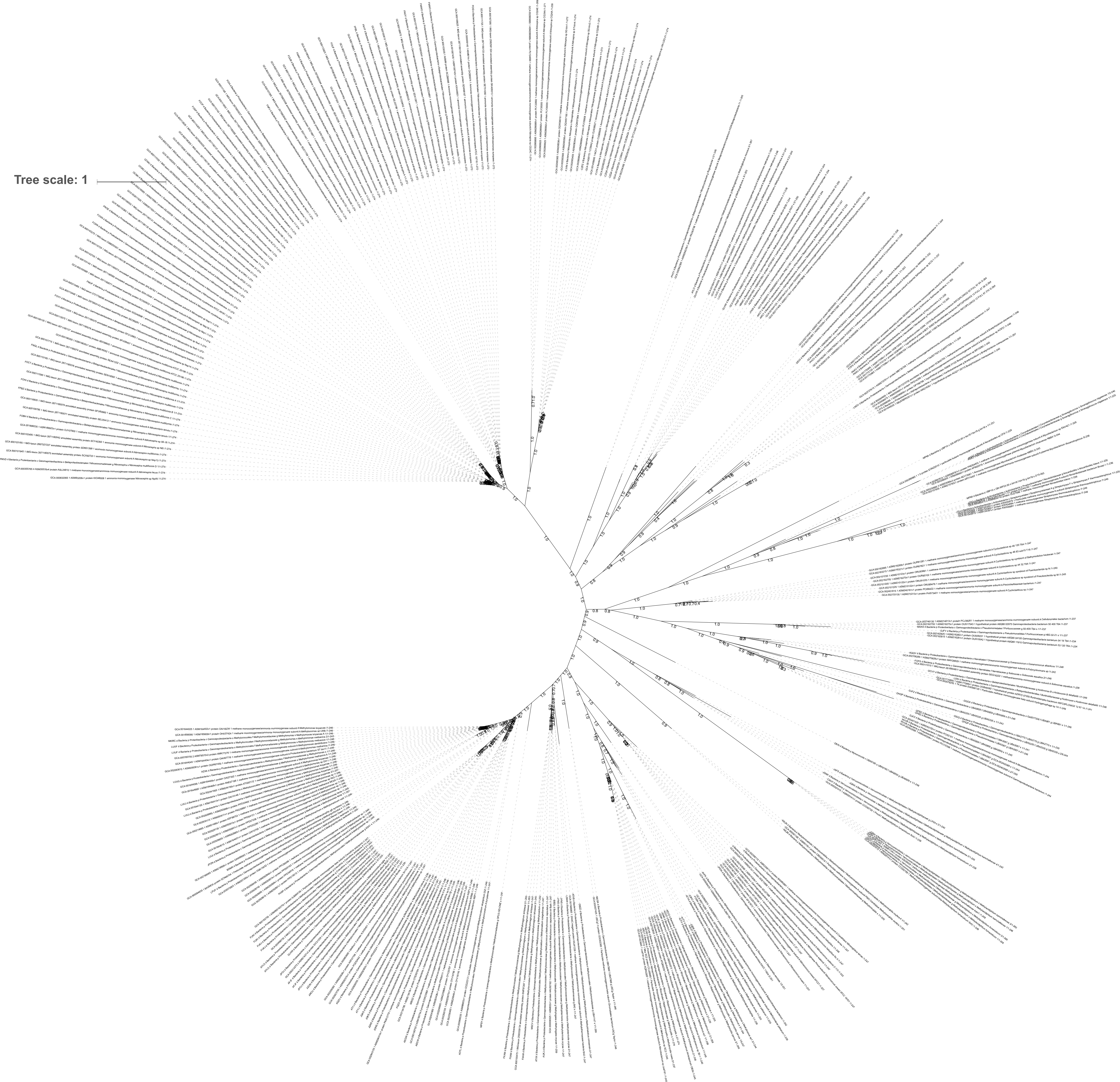

### Supplemental Figure 2

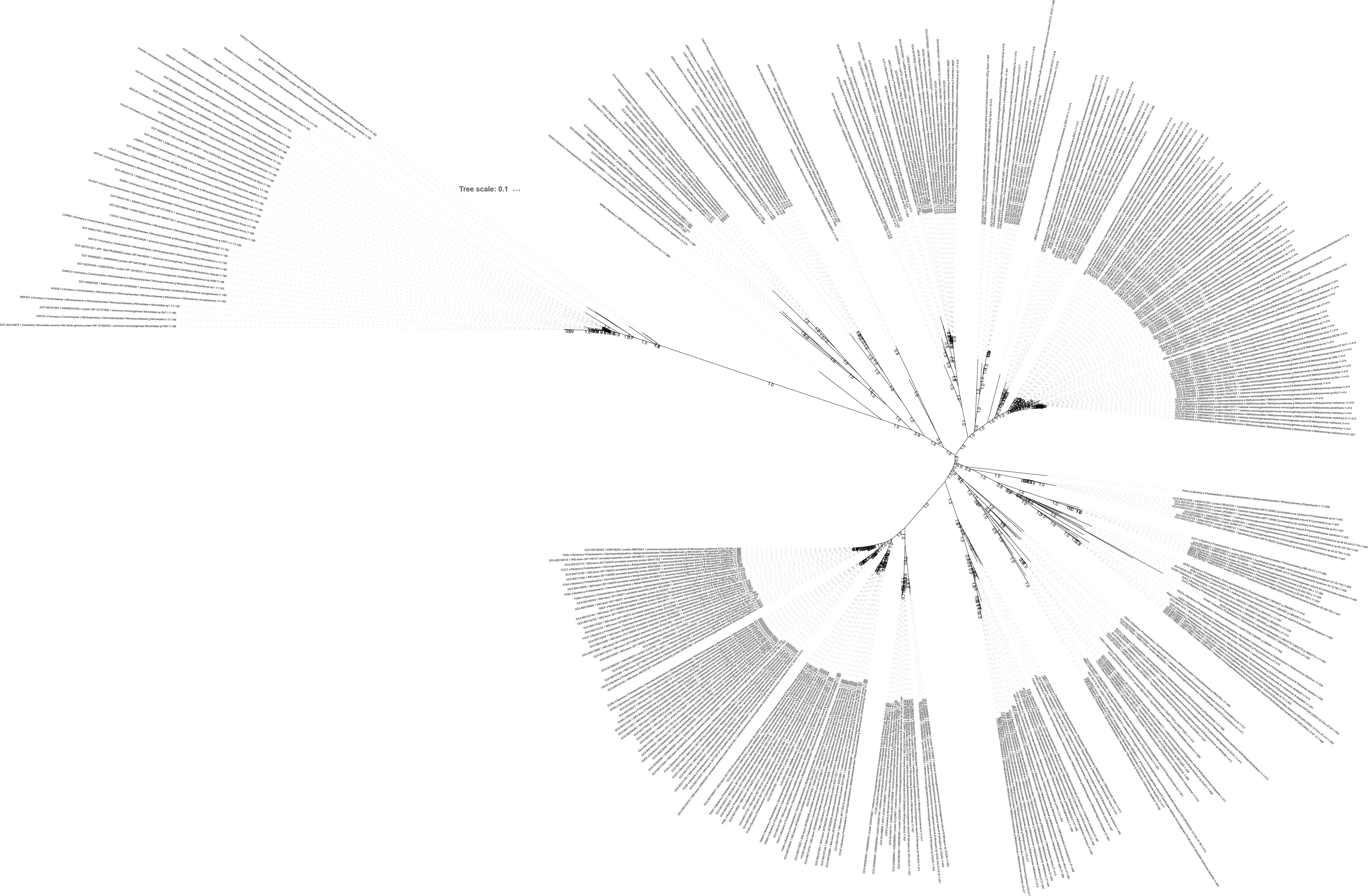

### Supplemental Figure 3

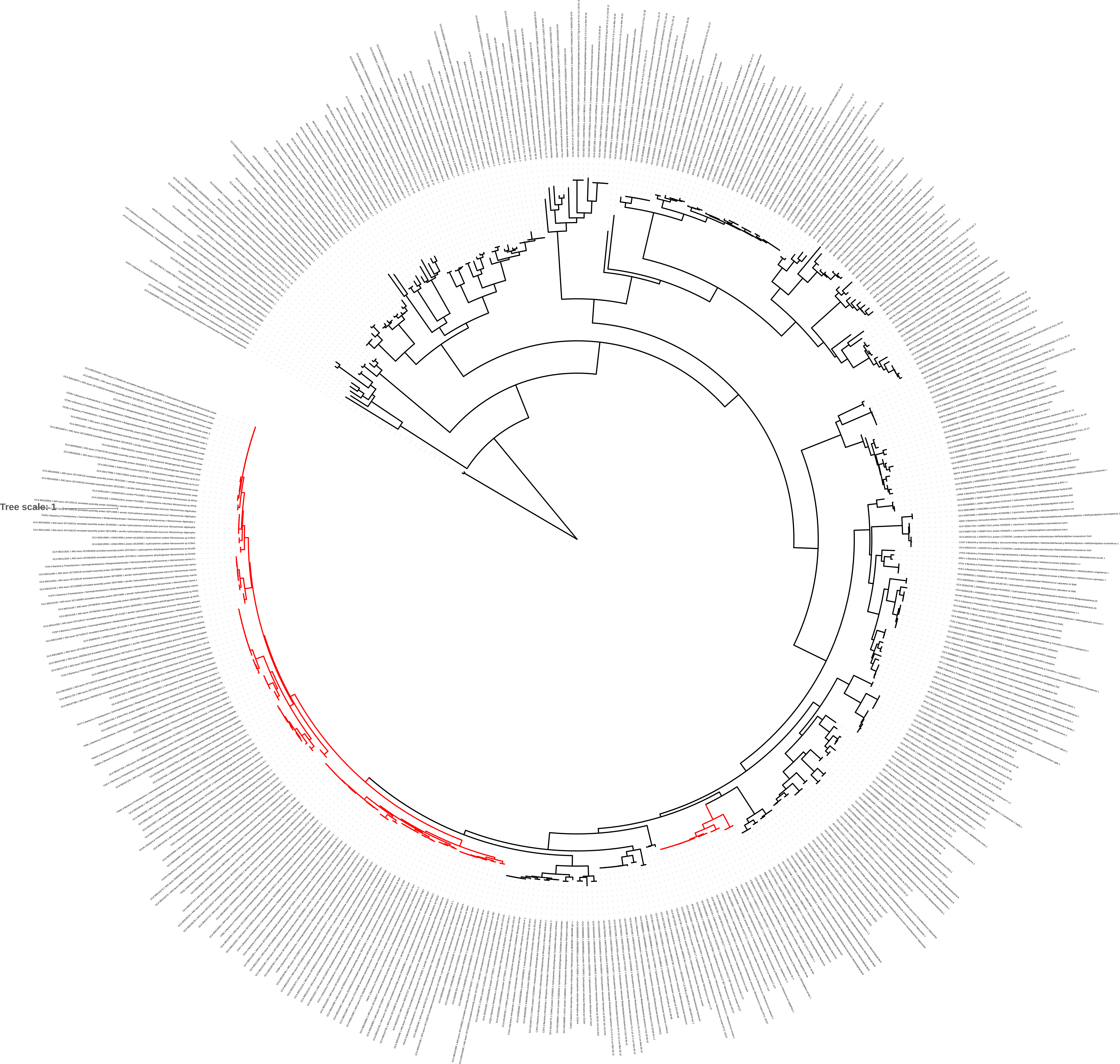

### Supplemental Figure 4

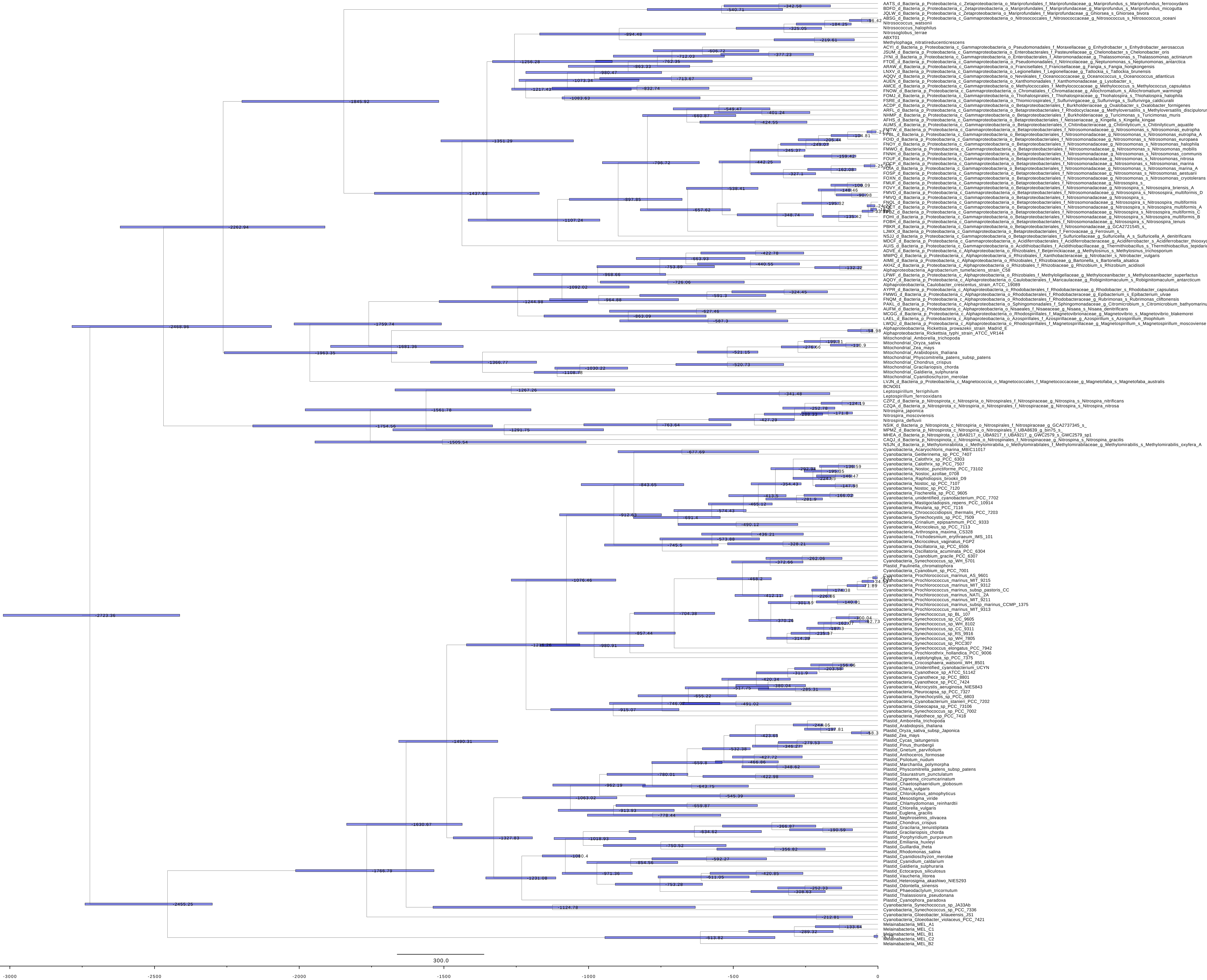
